# Metabolic mode estimated by breathing reflects long-term motor memory

**DOI:** 10.1101/2024.06.27.600911

**Authors:** Takuji Hayashi, Nobuyasu Nakano, Sohei Washino, Akihiko Murai

**Author notes:** Corresponding author. **Correspondence** Takuji Hayashi, The University of Tokyo, Graduate School of Education, Division of Physical and Health Education. **Author contributions** T.H., N.N., S.W., and A.M. designed the experiments. T.H. and N.N. performed the experiments. T.H. analyzed the data. T.H., N.N., S.W., and A.M. wrote the paper. **Competing interests** The authors declare no competing interests.

## Abstract

Respiration is a crucial metabolic process that converts macronutrients, carbohydrates and fats, and oxygen (O2) into energy and carbon dioxide (CO2) to support motor actions. In addition to the energy demands of movements, the brain is a significant energy consumer, accounting for approximately 20% of the body’s total energy expenditure and relying primarily on carbohydrates for neural activity and plasticity. However, it is not known whether O2-CO2 gas composition in breathing can serve as an indicator of neural activity and plasticity as they can for movement intensity. In the human reaching movement tasks, we evaluated time-constants of sensorimotor learning while recording O2-CO2 gas exchange. We computed the respiratory exchange ratio (RER), indicating which carbohydrate or fat is used preferentially, and found that the RER was unaffected by the execution and learning of reaching movements and that it was stable within individuals but varied across individuals. Interestingly, using computational modeling to identify short and long-time constants of sensorimotor learning, individual RER levels correlated with the magnitude of long-term, but not short-term memory. Furthermore, to experimentally manipulate the individual RER, we provided 200 kcal of glucose immediately before the task. Surprisingly, this simple intervention dramatically increased 24-hour retention by 21%. Together, the RER served as a remarkable proxy for long-term motor memory, and glucose intake shifted the physiological idling state for sensorimotor learning.

## INTRODUCTION

Respiration is a crucial process in metabolism responsible for the generation of adenosine triphosphate (ATP), an energy currency that supports all motor actions (Fig. 1a). Metabolism converts macronutrients, primarily carbohydrates and fats, along with oxygen (O2), into ATP and carbon dioxide (CO2). These metabolic processes exhibit varying rates and efficiencies. Quick, high-intensity motor actions require rapid energy production from carbohydrates, whereas prolonged, low-intensity motor actions require significant energy production from fat. Several studies have shown that the intensity of movement, measured by force or muscle activity, can be approximated with gas composition in breathing and is energetically optimized by motor experience (Abram et al., 2022; Huang et al., 2012; Selinger et al., 2015; Shadmehr et al., 2016). Therefore, breathing is recognized as a reliable measure for estimating the energy expenditure associated with movement execution.

**Figure 1.**
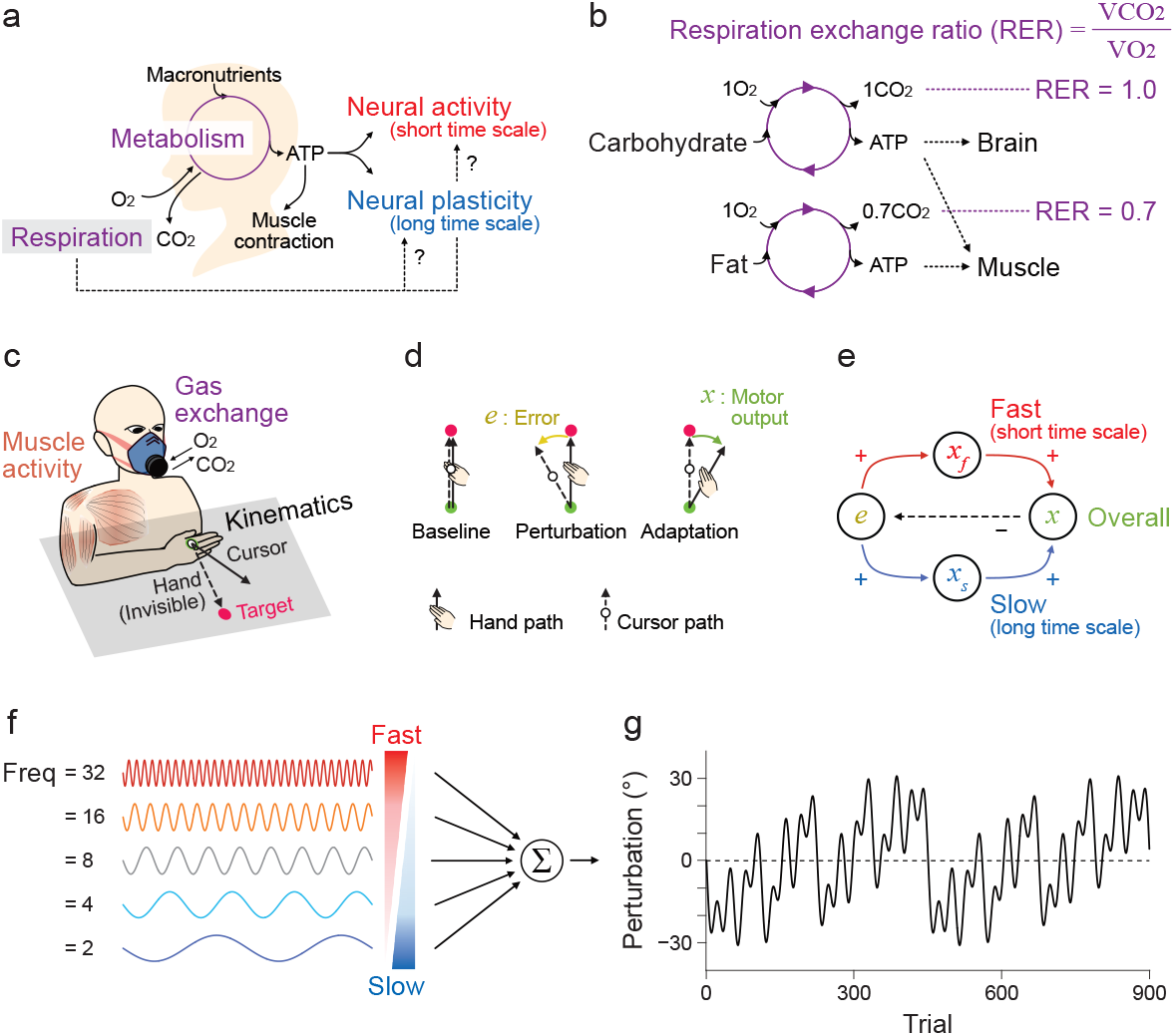
Respiratory exchange ratio and its potential role for sensorimotor learning. a. The metabolic cycle converts oxygen (O2) and macronutrients, carbohydrates, and fats, into carbon dioxide (CO2) and adenosine triphosphate (ATP), the energy currency that supports neural activity, plasticity, and muscle activity. Breathing, which involves O2-CO2 exchange, plays an important role in this process, but it is unknown whether breathing can predict learning and memory, which are influenced by neural activity and plasticity. b. The respiratory exchange ratio (RER), calculated as the ratio of CO2 production (VCO2) to O2 consumption (VO2), reflects the metabolic preference for using carbohydrates or fats as fuel. In the cortex, where only carbohydrates are used, the RER approaches 1.0. c. We designed Experiment 1 to test whether the RER, calculated by O2-CO2 exchange, represents sensorimotor learning of reaching movements while recording muscle activity. d. Visuomotor rotation (VMR), which angularly shifts the position of the visual cursor from the hand position, was introduced to elicit sensorimotor learning. This example shows counterclockwise VMR; the leftward visual error shifts the reaching movement to the right. e. A theory posits that two motor memories with different time constants, fast and slow, are stored in sensorimotor learning. The short- or long-time constants of sensorimotor learning, which presumably correspond to neural activity and plasticity. f. To accurately estimate motor memories with varying time constants, a dynamic VMR comprising five sinusoidal functions with different frequencies ranging from 2 to 32 Hz. It is hypothesized that rapidly changing VMR induces fast motor memory formation, while gradually changing VMR elicits the development of slow motor memory. g. The sum of the sinusoidal function is used in the reaching movement task.

In addition to the energy demands for muscle contraction, the brain is a significant energy consumer, utilizing 20% of the total energy expenditure despite comprising only 2% of body weight (Raichle and Gusnard, 2002). In particular, cortical energy is crucial for both neural activity (Oakes et al., 2004; Sokoloff, 1977) and plasticity (Lee et al., 2013; zur Nedden et al., 2011), which must play pivotal roles for sensorimotor learning. Sensorimotor learning not only establishes new patterns of neural activity (Li et al., 2001; Paz et al., 2003; Sun et al., 2022) but also induces novel cortical connections through long-term potentiation-like mechanisms (Kida et al., 2016; Landi et al., 2011; Rioult-Pedotti et al., 2000, 1998). Notably, these processes operate on different time scales: trial-by-trial emergence of new neural activity (Perich et al., 2018; Veuthey et al., 2020) contrasts with the slower, structural consolidation of connections, such as dendritic spine growth within an hour at the earliest (Harms et al., 2008; Xu et al., 2009). This temporal dichotomy may mirror a prevalent computational framework in sensorimotor learning, in which fast and slow processes interact to update motor memory (Smith et al., 2006). The fast process adapts quickly but decays rapidly, whereas the slow process learns gradually and supports longer-term retention. Computational models incorporating these dynamics successfully capture behavioral features of both short- and long-term memory formation (Joiner and Smith, 2008). Given the energetic demands of both fast and slow learning processes, we hypothesized that breathing metrics—particularly those reflecting cerebral metabolism—might provide a window into the temporal characteristics of sensorimotor learning.

To test this hypothesis, we conducted human behavioral experiments involving reaching movement tasks while recording O2-CO2 gas exchange. We focused on the respiratory exchange ratio (RER, Fig. 1b), which is the volume of CO2 produced divided by the volume of O2 consumed (McArdle et al., 2010), as it indicates the metabolic mode. Carbohydrate metabolism converts O2 to CO2 in a 1:1 ratio (RER = 1.0), while fat metabolism converts O2 to CO2 in a 1:0.7 ratio (RER = 0.7). Thus, the RER theoretically varies between 0.7-1.0, depending on the macronutrient usage. Since the brain utilizes only carbohydrates for the energy resource, its RER in the cortex is 0.97-0.99 (Himwich and Nahum, 1932), implying that a higher RER could signal increased cerebral energy expenditure. We observed that the RER was not influenced by the execution and learning of reaching movements; rather, it was stable within individuals but varied across individuals. Interestingly, using a computational decomposition of motor memory into fast and slow components, we discovered that individual RER levels correlated with the magnitude of slow motor memory, but not fast memory. Furthermore, we provided 200 kcal of glucose immediately before the experiment to test the hypothesis that manipulation of the RER can modulate long-term motor memory. This simple intervention increased the RER on day 1, and surprisingly, enhanced motor memory retention by 21% on day 2. Together, these findings suggest that the RER can serve as a physiological proxy for long-term motor memory, and that modulating metabolic state with glucose intake before learning can shift the energetic baseline, thereby enhancing memory consolidation.

## RESULTS

### Sensorimotor learning of dynamic perturbation in Experiment 1

Experiment 1 aimed to distinguish the short- and long-time constants of sensorimotor learning while monitoring O2-CO2 gas exchange and muscle activity in 18 human participants (Fig. 1c, 18 males aged 21-52 years). Participants engaged in planar reaching movements controlling a visual cursor on a display, where a visuomotor rotation (VMR) shifted the visual cursor away from hand movements angularly (Fig. 1d). To dissect the time constant of sensorimotor learning (Fig. 1e), we introduced a dynamic VMR perturbation (Fig. 1f and g), comprising 10°-amplitude sinusoidal sequences of VMR with different time constants ([2, 4, 8, 16, and 32] cycles over 900 trials) (Miyamoto et al., 2020). The two-state model posits that, in sensorimotor learning, two motor memories are updated concurrently with different time constants (Smith et al., 2006). This unique idea, other from more recent frameworks (Heald et al., 2021), is leveraged to reliably estimate time features of motor memory, where slow motor memory predicts next-day retention at 99% of variances in the group-level analysis and at 50% of variances in the individual level (Joiner and Smith, 2008). Accordingly, we predicted that fast motor memory would adapt to VMR changes more quickly (16 and 32 cycles), while the slow motor memory would adapt to VMR changes more gradually (2 and 4 cycles). The dynamic VMR allowed for an adequate assessment of learning components with different time constants.

Overall movement directions largely followed changes in the dynamic VMR (Fig. 2a). Fourier transform analysis revealed large amplitudes only at trained frequencies ([2, 4, 8, 16, and 32] Hz), indicating appropriate adaption to the dynamic VMR (Fig. 2b). Despite not reaching the perturbation levels of 10°, the mean amplitudes at the trained frequencies (6.97° ± 0.29°, mean ± SEM) significantly surpassed those at the untrained frequencies between 2 and 32 Hz (0.56° ± 0.10°) (Fig. 2c, t(17) = 19.99 and p = 3.0 × 10^−13^). These results indicate appropriate participant adjustment to the dynamic VMR.

**Figure 2.**
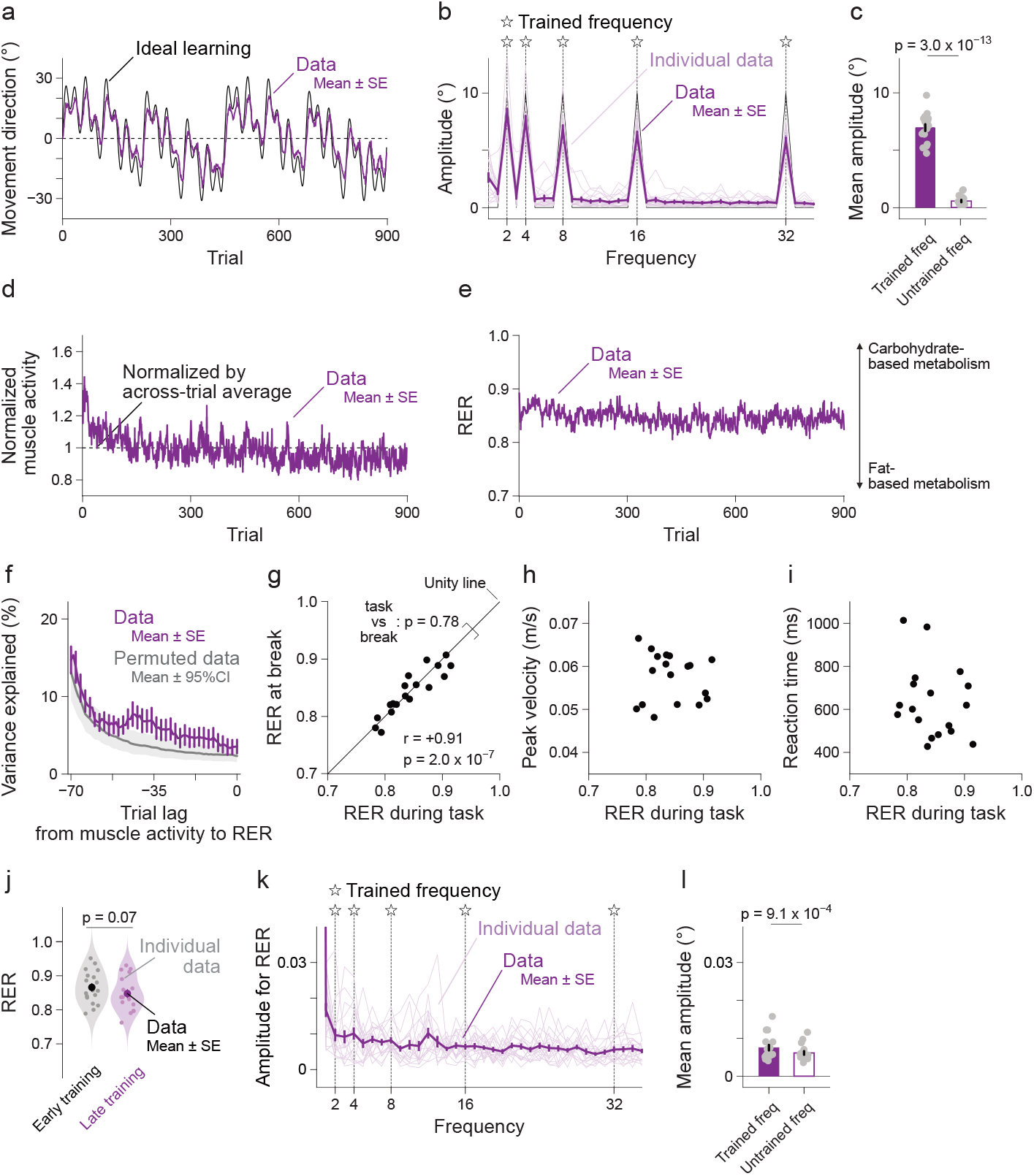
Few effects of execution and learning of reaching movements on RER. a. Movement directions changed with the VMR. Error bars show the standard errors of their means. b. Significant peaks at the trained frequencies were found with the Fourier transformation for a series of movement directions. c. The average of the peaks was significantly larger than that at the untrained frequencies. d,e. Self-normalized muscle activity (d) was high at the beginning of the experiment and decreased rapidly over the first 300 trials. In contrast, RERs (e) were very stable over the course of the experiment. f,g. Regression analysis with trial lags revealed that self-normalized muscle activity cannot explain the changes in RER (f, permutation test, p > 0.05 for all lags with FDR correction). Furthermore, there was no change between task and break (g). h,i. No relationships with possible factors, peak velocity (h) and reaction time (i), were observed. j. No differences between RER in the first and last block; Sensorimotor learning did not modulate RER. k,l. No obvious peaks at the trained frequency were found with the Fourier transformation for a series of RER (k), although the mean amplitude was statistically higher at the trained frequency (l).

### Effects of reaching movements on RER

The RER is known to covary with the intensity of movement execution (Goedecke et al., 2000); therefore, changes in muscle activity could potentially confound the relationship between the RER and sensorimotor learning. Thus, we investigated how sensorimotor learning altered muscle activity and whether these changes affected the RER. Electromyography (EMG) data from representative muscles including the pectoralis major, posterior deltoid, biceps brachii, and triceps brachii were recorded to observe movement execution intensity effects. Muscle activity normalized by the trial average was high at the experiment’s outset but quickly decreased, stabilizing at approximately one-third of the experiment’s duration (Fig. 2d). This pattern, recognized as the relaxation of co-contraction into a novel environment (Franklin *et al*., 2003), contrasted with the stability observed in the mean RER across participants, which did not fluctuate across trials (Fig. 2e), suggesting a deviation from typical sensorimotor learning patterns (Fig. 2a).

We examined whether muscle activity could account for RER variance across trials. Performing a linear regression with trial lags, we found that only 3.6% of the variance could be explained without lags, with minimal improvement observed across trials, even when accounting for trial lags (Fig. 2f). This was in comparison with the permuted data from randomized trial blocks without significance noted for all trial lags after false discovery rate (FDR) correction (p > 0.05). A slight enhancement was detected with 36-46 trial lags (p < 0.05, without FDR correction). However, these trials elapsed approximately 190-243 seconds (trial duration was 5.3 ± 0.1 seconds), which is 1.5-2.0 times longer considering the time constant of the metabolic loop compared with the previous report showing a 120-second lag (Short and Sedlock, 1997).

To further explore the impact of movement execution, we compared the average RER during the reaching movement tasks and breaks (Fig. 2g). Despite high consistency within participants (r = +0.91 and p = 2.0 × 10^−7^), the absence of a statistically significant difference between task and break RERs (t(17) = 0.29 and p = 0.78) indicated minimal effect of movement execution. Additionally, RERs during tasks exhibited no correlation with mean peak velocity (Fig. 2h, r = −0.10 and p = 0.69) or reaction time (Fig. 2i, r = −0.26 and p = 0.30). These results suggest minimal influence of movement execution on the RER owing to the insufficient intensity of reaching movements.

### Effects of sensorimotor learning on RER

Next, we investigated whether there was any indication of sensorimotor learning reflected in the RER (Fig. 2e). However, we found that the RER remained constant during sensorimotor learning when comparing the RER between the early and late training blocks (Fig. 2j, t(17) = 1.91 and p = 0.07). Nonetheless, these values were highly consistent within participants (r = +0.65 and p = 0.0033).

Given that we used the dynamic VMR as the sum of the sinusoidal function, the amount of VMR to be learned in the early training block was equal to that in the late training block. Consequently, if the RER were to fluctuate with the amount of motor memory, it should remain stable during sensorimotor learning. In such a scenario, the peaks in the frequency domain of the RER are the same as those in the dynamic VMR. Alternatively, the RER might fluctuate with changes in motor memory, particularly following the derivatives of the sum of the sinusoidal functions, which would manifest as cosinusoidal function at the same frequencies. Thus, we would expect to observe peaks at [2, 4, 8, 16, and 32] Hz if sensorimotor learning affected the RER. Figure 2k illustrates the amplitudes of the frequency analysis, revealing no discernible peaks. However, we found a slight increase in the mean amplitudes at the trained frequencies compared to those at the untrained frequencies (Fig. 2l, t(17) = 4.01 and p = 9.1 × 10^−4^).

The significant difference in the RER at the trained and untrained frequencies may not necessarily signal sensorimotor learning for the following three reasons. First, the absolute difference (0.0017) was smaller than the standard deviation of the within-participant RER fluctuation (SD: 0.044 ± 0.003). Second, the RER may not behave as a random process as white noise, it may exhibit characteristics of pink noise (or 1/f noise), where the power is inversely proportional to the frequency. Such patterns of pink noise are widely found in human physiology (Pritchard, 1992; Zarahn et al., 1997) and cognition (Gilden et al., 1995), and the RER’s tendency to display larger amplitudes at low frequencies (Fig. 2k) could imply a pattern of pink noise. If the RER is 1/f scaled, the mean amplitudes at the trained frequencies will be 2.41 times larger than those at the untrained frequencies. Third, the Fourier analysis could not identify polarity, meaning that we cannot ascertain whether the RER increased or decreased with sensorimotor learning. To further investigate whether the RER changed in tandem with sensorimotor learning, we analyzed RER in Experiment 2 (see below).

### Individual variability in RER reflects slow motor memory

We designed the dynamic VMR to rigorously estimate both fast and slow components of sensorimotor learning. Individual data were fit into a two-state model. The overall motor output followed this behavior (Fig. 3a, R^2^ = 0.82 ± 0.04). As predicted, we observed peaks at the trained frequencies for both fast and slow components, as well as overall (Fig. 3b). The mean amplitude at the trained frequencies was significantly higher for the overall data (Fig. 3c, t(17) = 24.17 and p = 1.3 × 10^−14^). However, neither the experimental data nor the overall data correlated with the RER during the task (Fig. 3d, r = 0.04 and p = 0.88 for experimental data, r = 0.16 and p = 0.53 for overall), suggesting that the RER does not reflect overall sensorimotor learning.

**Figure 3.**
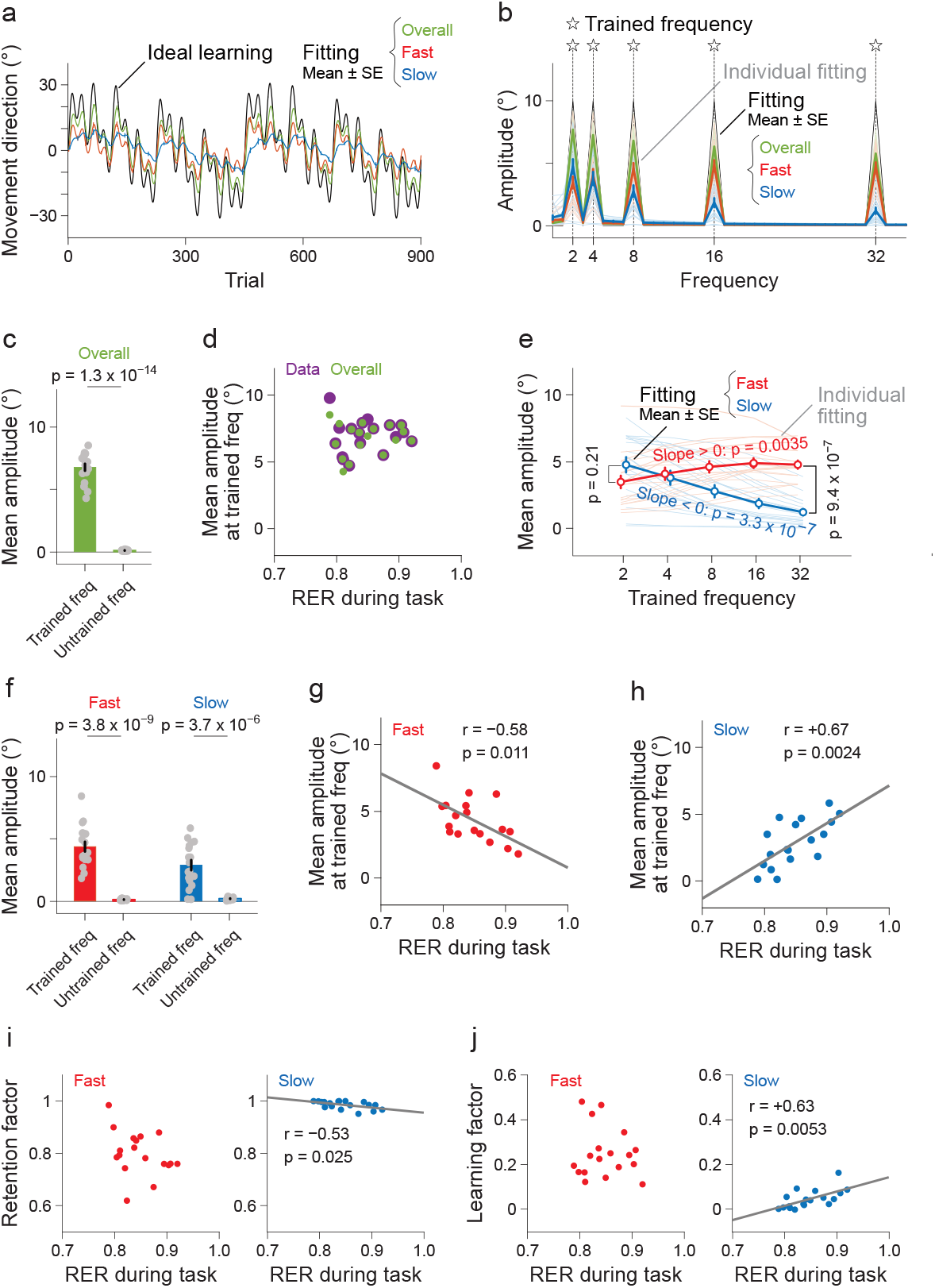
Individual RER reflects slow components of sensorimotor learning. a-c. The two-state model captured the sensorimotor learning processes (a). Obvious peaks of the overall, fast, and slow components at the trained frequency were revealed by the Fourier transformation (b), and the mean amplitudes at the trained frequency were higher than those at the untrained frequency (c), analogous to the experimental data shown in Figure 2c. d. The experimental and fitted data were not correlated with the RER during the task. e,f. The amplitudes of the fast and slow components were increased and decreased, respectively, with the trained frequency (e). The mean amplitudes at the trained frequency were higher than those at the untrained frequency (f). g,h. The mean amplitudes of the fast (g) and slow (h) components were negatively and positively correlated with the RER during the task, respectively. i,j. For comparison with the model parameters, the RER during the task was correlated only with the slow retention factor (i) and the slow learning factor (j).

Figure 3e illustrates that the peak amplitudes for the fast components gradually increased with frequency (slope > 0, t(17) = 3.39 and p = 0.0035), while those for the slow components gradually decreased (slope < 0, t(17) = 8.07 and p = 3.3 × 10^−7^). A statistically significant difference between fast and slow components was found at the highest frequency of 32 Hz (t(17) = 7.45 and p = 9.4 × 10^−7^), with a marginal difference at the lowest frequency of 2 Hz (t(17) = 1.32 and p = 0.21). The mean amplitudes at the trained frequency were higher than those at the untrained frequency (Fig. 3f, t(17) = 10.98 and p = 3.8 × 10^−9^ for the fast component, t(17) = 6.71 and p = 3.7 × 10^−6^ for the slow component). Interestingly, a statistically significant negative correlation was found for the fast component (Fig. 3g, r = −0.58 and p = 0.011), but a positive correlation was found for the slow component (Fig. 3h, r = 0.67 and p = 0.0024) with the RER during the task. Note that only the slow component and the RER before the experiment were significantly correlated (r = −0.34 and p = 0.17 for the fast component, r = 0.47 and p = 0.048 for the slow component), suggesting that RER can predict the slow motor memory to be learned. However, this correlation was weaker than during the task (Fig. 3h), possibly owing to the time at baseline being slightly too far from the training phase.

Fast and slow motor memories were negatively correlated (r = −0.71 and p = 8.6 × 10^−4^) because an increase in one memory decreased the other, suggesting that the opposite correlation shown in Figure 3g and h would not satisfy the assumption that the RER reflects both of them. Therefore, we further examined whether the RER reflected the fitting parameters for the fast and slow, namely the retention and learning factors (Fig. 3i and j). Interestingly, we found a negative correlation with the retention factor (r = −0.53 and p = 0.025) and a positive correlation with the learning factor (r = 0.63 and p = 0.0053) of the slow component. However, no significant correlations were found for the fast component (r = −0.35 and p = 0.15 for retention factor, r = −0.14 and p = 0.59 for learning factor). These findings suggested that participants with a higher RER demonstrated faster learning but also exhibited greater forgetting in slow motor memory. Given that both changes in hand motion away from and toward the 0° target direction necessitate the updating of slow motor memory in the dynamic VMR, the results suggest that the slow component of sensorimotor learning is more flexible. The RER does not alter the execution and learning of reaching movements; instead, it reflects the slow component of sensorimotor learning, characterized by a motor memory with a relatively longer time constant.

### Long-term retention with glucose intervention: Experiment 2

We found that the RER reflected the slow component of sensorimotor learning in Experiment 1. The slow component serves as an estimate, offering indirect means to measure long-term memory. To directly investigate whether the RER reflects long-term memory, we conducted a test for 24-hour retention, wherein the slow components played a significant role (Fig. 4a) (Joiner and Smith, 2008). Furthermore, while the RER remained constant in Experiment 1, several methods have been proposed to alter it, including circadian rhythm (Zitting et al., 2018), dietary factors, and genetic predispositions (Toubro et al., 1998). Since glucose intake is a simple but powerful way to increase the RER (Tellez et al., 2013), we tested whether glucose intervention resulted in improved long-term motor memory (Fig. 4a).

**Figure 4.**
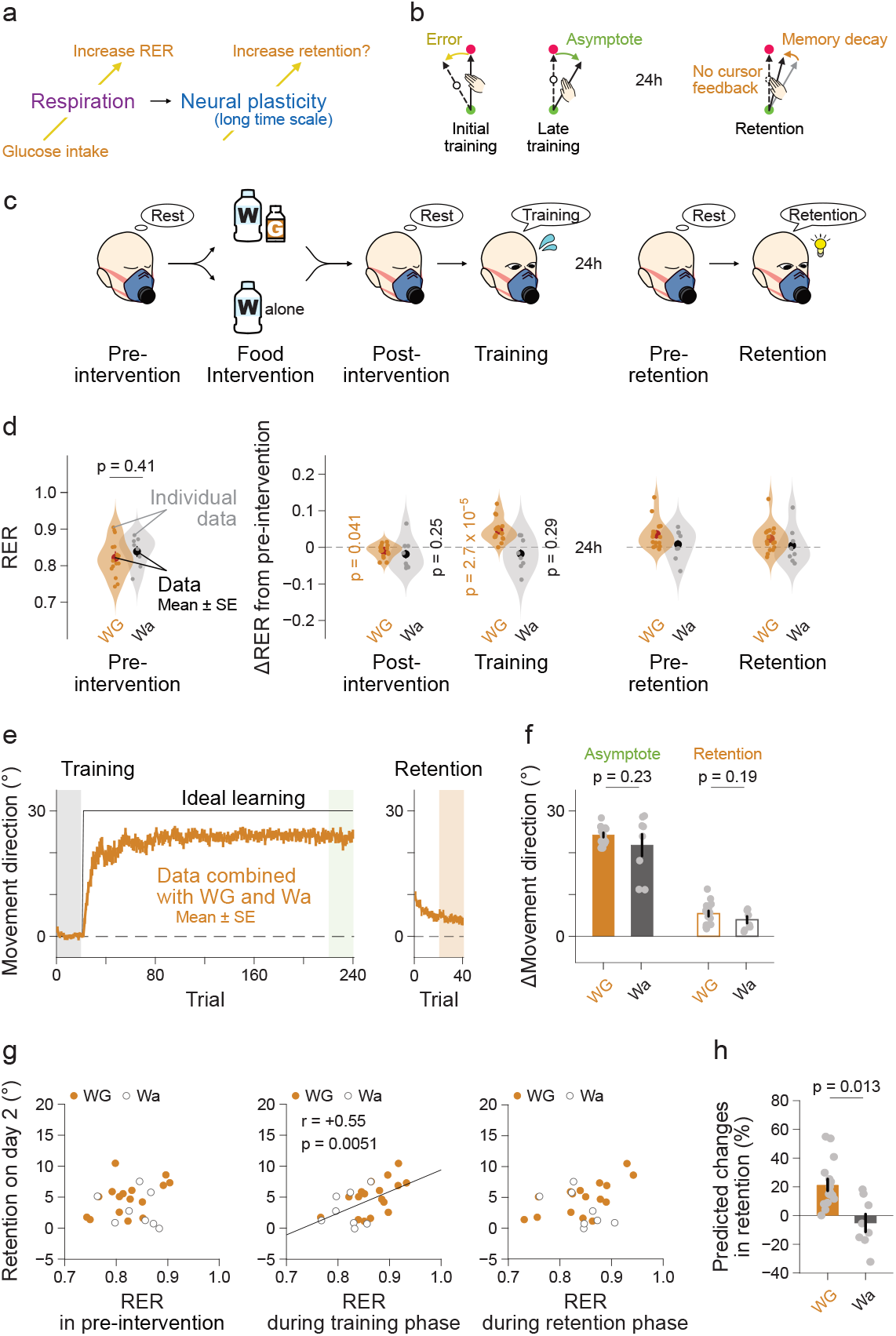
RER predicts glucose-intake enhancement of long-term motor memory. a. Based on the results of Experiment 1, which showed that the RER reflects slow motor memory, we tested the hypothesis that increasing the RER with glucose intake would improve 24-hour retention. b. Experiment 2 was designed to examine how much of the 30° constant VMR learned on day 1 was retained on day 2 during reaching movements without visual feedback. c. On day 1, the RER was recorded for 20 min before and after an intervention in which one group received 200 kcal glucose (WG) while the other group did not (Wa). Both then performed the same training task. On day 2, the RER was recorded for 20 minutes, followed by the retention task. d. No difference in the original RER was found in the pre-intervention phase. Furthermore, the change in the RER was not found in the post-intervention phase but in the training phase. The effects did not persist on day 2; no differences were found in the pre-retention and retention phases. e,f. We observed the typical learning curve for VMR training on day 1 (left panel in e). On day 2, the amount of learning largely dropped, but we can still see significant retention (right panel in e). Notably, no differences were found between the groups (f). g. The correlation analysis showed a significant positive relationship between retention and intervention RER, but no relationships against baseline RER or RER on retention. h. Changes in the RER from the pre-intervention phase to the training phase estimated a 21.4% ± 4.1% increase in retention with the glucose intervention.

We recruited an additional 24 human participants (14 males and 10 females aged 19-39 years) for Experiment 2. This was a two-day experiment in which they adapted to a constant VMR on day 1 and tested retention on day 2 while monitoring the RER (Fig. 4b). The participants were divided into two groups: water and glucose (WG, n = 16) and water alone (Wa, n = 8) (Fig. 4c). On day 1, the WG group consumed a gel-type drink containing 50 g glucose (200 kcal) and water, whereas the Wa group consumed only water. Before and after the intervention, resting-state gas exchange was recorded for 20 min to observe the rapid effects of glucose intervention on the RER. After a short baseline of 20 trials, they learned a constant VMR for 220 trials while the gas exchange was consistently recorded. On day 2, the WG and Wa groups returned to the laboratory, and we examined 20 min of resting-state gas exchange, followed by the retention test without visual feedback (Fig. 4b), which can measure long-term memory but not learning, while the gas exchange was recorded.

### Increase in RER following glucose intake

We found that the RER increased only after glucose intake. The RER before the intervention was not significantly different between the WG and Wa groups (Fig. 4d, t(22) = 0.85 and p = 0.41). The RER immediately after the intervention did not increase but rather slightly decreased from the pre- to post-intervention phases (t(15) = 2.23 and p = 0.041 for WG, t(7) = 1.24 and p = 0.25 for Wa) and was not significantly different between these groups (t(22) = 0.42 and p = 0.68). Notably, however, we found a significant increase in the RER from the pre-intervention to task phases only in the WG group (t(15) = 5.94 and p = 2.7 × 10^−5^ for WG, t(7) = 1.15 and p = 0.29 for Wa), with a significant difference noted between these groups (t(22) = 2.77 and p = 0.011). The increase in the RER did not persist for the next day, and no significant differences were found between these groups in either the pre-retention (t(22) = 0.43, p = 0.67) or retention phases (t(22) = 0.07 and p = 0.95). This result suggests that the RER can be successfully controlled by glucose intervention. Notably, there were no changes in the RER during the training phase in the Wa group, providing important evidence that sensorimotor learning does not affect the RER.

### Association between individual RER responses to glucose intake and retention

We observed basic trends in the VMR learning curves. After short baseline periods, the hand movement directions gradually shifted toward ideal learning (Fig. 4e). The resulting learning levels were 24.2° ± 0.6° for the WG group and 21.8° ± 2.6° for the Wa group, which were not significantly different (t(22) = 1.25 and p = 0.23). The learning amounts at the retention period on day 2 were obviously reduced from the end of the training period, but were still greater than the baseline at 5.5° ± 0.7° for the WG group but 3.9° ± 0.8° for the Wa group. Unfortunately, the retention amounts were not significantly different between groups (t(22) = 1.36 and p = 0.19). Because the RER in the training phase was so variable (Fig. 4d), a comparison at the group level would not reveal the effects of glucose intake on retention.

The results of Experiment 1 suggested that RER during the task reflected slow motor memory (Fig. 3h). To provide a consistent analysis, we performed a correlation of retention on day 2 with RER at three representative phases: in the pre-intervention, during the task on day 1, and during the task on day 2. We found no significant correlation between the retention and the RER in the pre-intervention phase (r = 0.16 and p = 0.45) or during the task on day 2 (r = 0.31 and p = 0.14), but a significant correlation was found with the RER during the task on day 1 (r = 0.55 and p = 0.0051) (Fig. 4g). The results indicated that the original RER that the participants inherently had in the pre-intervention phase did not predict retention, but the intervention RER that increased with glucose intake in the training phase predicted retention, suggesting that the control of RER via glucose intake can increase retention at the intra-individual level.

How much can retention be improved by a 200 kcal glucose intake? Using the regression parameters, slope and offset, from the RER during the task on day 1 to retention on day 2 (Fig. 2g), we predicted the retention levels if the participants inherently had the original RER during training. We also projected the RER during the task on day 1 onto the regression line and calculated the increase in RER within individuals (See Methods section). Surprisingly, the glucose intake improved up to 21.4% ± 4.1% in the WG group (t(15) = 5.25 and p = 9.8 × 10^−5^), but the water alone did not improve at −5.1% ± 6.2% in the Wa group (t(7) = 0.83 and p = 0.43) (t(22) = 3.67 and p = 0.013 between the two). The results demonstrate that a simple glucose intervention improves long-term motor memory acquisition.

## DISCUSSION

Respiration plays a crucial role in energy production and supports neural activity and plasticity (Lee et al., 2013; Oakes et al., 2004; Sokoloff, 1977; zur Nedden et al., 2011), which are essential for sensorimotor learning. Breathing patterns provide detailed information about whole-body metabolism including movement intensity (Abram et al., 2022; Huang et al., 2012; Selinger et al., 2015; Shadmehr et al., 2016); however, it is largely unknown whether they are also a prominent index of brain metabolism, that is, whether breathing patterns predict sensorimotor learning. According to the two-state sensorimotor learning model (Smith et al., 2006), the faster component reflects short-term memory, whereas the slower component reflects long-term memory. We investigated whether the metabolic mode estimated by the RER reflects short-term or long-term memory. The results showed that the RER was not influenced by the execution or learning of reaching movements (Fig. 2). Instead, the RER predicted the amount of estimated slow rather than fast motor memory (Fig. 3). We also investigated whether metabolic interventions could improve long-term memory, that is, next-day retention. To accomplish this, we provided 200 kcal of glucose, which immediately shifted the participants towards more carbohydrate utilization. Surprisingly, this intervention not only increased the RER but also improved next-day retention by up to 21% (Fig. 4). Together, with two separate cohorts of human participants, these findings showed that the metabolic mode estimated by the RER can serve as an indicator of long-term motor memory, particularly at the intra-individual level, implying a potential avenue for enhancing human motor performance.

### Methodological consideration

The metabolic mode estimated by the RER varied widely from 0.74 to 0.97 among individuals even at baseline, which is very similar range to the previous findings (e.g., from 0.718 to 0.927 in (Goedecke et al., 2000)). Previous studies have shown that during low-intensity activities such as unloaded or lightly loaded pedaling—activities likely to exert more load than the reaching movements in our study—RER does not significantly deviate from baseline levels (Wasserman et al., 1973). This is consistent with our findings (Fig. 2e). We also found no explanation of muscle activity on RER (Fig. 2f) as well as unchanged RER between at-task vs at-break (Fig. 2g). Furthermore, the RER is highly stable even during sensorimotor learning (Figs. 2e and 4d gray) and no correlation against sensorimotor learning amplitudes (Fig. 3d). The results suggest that the execution and learning of reaching movement do not affect RER accordingly.

We believe that a comparison between the presence and absence of glucose intake provides a direct examination of how the RER intervention affects long-term motor memory. However, the baseline RER variability was high (SD across participants: 0.049) compared to the increased RER (0.046 ± 0.008 from pre-intervention to task phases) with only 200 kcal of glucose intake, implying that the glucose intervention on next-day retention (Fig. 4f) would be buried under large individual differences at baseline. The long-term motor memory would be enhanced by maximizing the physiological effects of glucose intake, such as the glucose dose or glucose digesting time. Notably, a literature review showed enhancement of memory with glucose intake is limited even in the other memory types, such as verbal word memory (Smith et al., 2011). Our results underscore a new type of memory retention improvement at the intra-individual level, and it needs to turn the tide for investigation of physiological basis for sensorimotor learning.

Glucose intake may contribute to other cognitive improvements during learning. For example, glucose intake may induce a positive mood and reduce hunger, perhaps leading to an upregulation of motivation and attention. These cognitive improvements would affect the fast component, reflecting the use of an explicit strategy (McDougle et al., 2015), rather than the slow component. Thus, if glucose intake elicited only cognitive improvements, the slow component would not change. Our findings that the glucose intake affects slow, long-term motor memory suggest that cognitive improvements from the glucose intake are not the main driver of its enhancement of long-term motor memory.

### The cortical metabolism and time-scale dynamics in sensorimotor Learning

Long-term motor memory is shaped by multiple factors, and our computational approach serves to simplify the intricate interplay of biological factors of sensorimotor learning, such as protein synthesis and gene expression (Kandel et al., 2014). For instance, neural plasticity is regulated by protein synthesis (Luft et al., 2004). The metabolic demands associated with sensorimotor learning may further influence neuroplasticity through mechanisms involving ATP availability and mitochondrial function. However, the specific connections between metabolic mode and processes of motor memory retention remain unclear and require further investigation.

Another possible explanation is cortical-area-dependent time constants (Kim et al., 2015). The parietal areas and cerebellum, which are considered as key regions in the neural basis of sensorimotor learning, have been suggested to exhibit neural activity changes at different time scales—fast for the parietal areas and slow for the cerebellum. A study by Vaishnavi and colleagues demonstrated that these areas also differ in their metabolic profiles, particularly in terms of aerobic glycolysis (Vaishnavi et al., 2010). Specifically, aerobic glycolysis is markedly elevated in the medial and lateral parietal cortices, while the cerebellum exhibits levels well below the brain average. These findings suggest a potential coupling between regional metabolic signatures and time-dependent learning processes, suggesting that metabolic measures may serve as physiological markers of engagement in distinct brain systems during sensorimotor learning. Future work should aim to integrate metabolic modeling with neural network dynamics to develop a more comprehensive picture of motor memory formation.

### The RER would indicate physiological idling state for sensorimotor learning

The brain accounts for 20% of total energy expenditure (Raichle and Gusnard, 2002). This proportion underscores that RER may indeed provide valuable insights into the neural states associated with learning and memory. However, it is important to note that the RER is an indicator of the net metabolic mode of the entire body. Although the contribution of brain metabolism is significant, it is certainly not a large contribution. One could argue that the effect of cortical metabolism on RER is limited. How does RER reflect long-term motor memory?

It should be noted that the glucose intake did not immediately increase the RER in the post-intervention phase, but rather slowly increased it during the subsequent training phase (Fig. 4d). This temporal pattern suggests a gradual physiological response that unfolds over time, that is, insulin secretion. Insulin secretion is not an instantaneous response but occurs gradually in response to increasing levels of glucose in the bloodstream. Insulin secretion is usually accompanied by an increase in RER (Dickson et al., 1924), and presumably regulates glucose levels in the body and neuronal and glial cells (Milstein and Ferris, 2021; Uemura and Greenlee, 2006) for future energy production. Thus, this increase in cerebral glucose levels would meet the cortical energy needs, especially for learning and memory.

Interestingly, insulin modulates memory formation, as evidenced by studies on Alzheimer’s disease, a model of impaired declarative memory function. Alzheimer’s disease is accompanied by disrupted glucose regulation and utilization (Kyrtata et al., 2021); however, surprisingly, insulin injection improves memory function in Alzheimer’s patients (Craft et al., 1996; Reger et al., 2008). Consistent results have been found in healthy subjects, indicating that the intranasal administration of insulin upregulates memory function (Benedict et al., 2007, 2004). Furthermore, insulin resistance in healthy obese individuals impairs their ability to learn; however, the administration of liraglutide, which temporarily stimulates insulin release, can restore this ability (Hanssen et al., 2023). Collectively, these studies suggest that insulin secretion and its receptors regulate memory. In line with these studies, our results suggest that the RER, which may be linked to insulin release, would reflect a physiological idling state for sensorimotor learning.

## METHODS

### Participants

In total, 18 participants (18 males, aged between 21 and 52 years) in Experiment 1 and 27 participants (16 males and 11 females, aged between 19 and 39 years) in Experiment 2 were recruited; however, we removed data from three participants in Experiment 2 owing to their data quality (See Data Analysis). All participants were right-handed, reported no neurophysiological disorders, and provided written informed consent prior to the experiments in accordance with the Declaration of Helsinki. The Committee on Ergonomic Experiments of the National Institute of Advanced Industrial Science and Technology (No. HF2022-1183) approved the protocol of Experiment 1 and the research ethics committee of the University of Tokyo (No. 23-204) approved the protocol for Experiments 1 and 2, respectively.

### Reaching movement task

Participants performed rapid 20-cm point-to-point arm reaching movements with their right hand while sitting on a chair positioned at the center of the task space. We used an exoskeleton robotic manipulandum in Experiment 1 (KINARM Exoskeleton Lab, Bkin Technologies, Canada), which can prevent unwanted fatigue and maintain a constant posture to reliably record muscle activity with electromyography (EMG), while we used a digitizing tablet and pen encased in a light grip in Experiment 2 (Wacom Intuos Pro L, Wacom, Japan) to reduce muscle activity that might affect the breathing data. The sampling rates were 1000 Hz in Experiment 1 and 200 Hz in Experiment 2, and the refresh rates of the visual screen were both 60 Hz.

At the beginning of each trial, the participants were asked to hold a white circle (1.0 cm in diameter) representing their hand position on a red start circle (1.5 cm). One second later, the red target circle (1.5 cm) was displayed 20 cm in front of the start circle. One second later, the target color changed to green, which was a cue to initiate a forward-reaching movement. Upon reaching the green target, the target color changed to blue and the start circle turned green, which was a cue to initiate a backward-reaching movement. Outback movements were completed, and the next trial started after an inter-trial interval of 1 s. A preliminary experiment showed that the outback movements took approximately 2 s. A trial duration of approximately 5 s was chosen because each trial would involve at least one breath, which occurred approximately once every 4 s. After the participants familiarized themselves with the task, we introduced VMR, which involved angularly shifting the cursor position.

To dissociate the fast and slow components of sensorimotor learning in Experiment 1, we designed the sum of sinusoidal functions(Miyamoto et al., 2020). The frequencies of the functions were [2, 4, 8, 16, 32] Hz over 900 trials; that is, the perturbation sequence combined gradual components (2 and 4 Hz) and sudden components (16 and 32 Hz). The amplitude of the function was 10°. The maximum amplitudes of the dynamic perturbation were ±30.9° and maximum changes in a trial were ±4.3°. We counterbalanced the signs of the amplitudes of the 18 participants. A total of 900 training trials were divided into 12 blocks, with an approximately one-minute break between blocks.

In Experiment 2, to test the 24-h retention with the glucose intervention, participants adapted to a constant VMR perturbation on day 1 and were tested for motor memory without visual feedback on day 2. The constant perturbation was 30°, which was counterbalanced across the 24 participants. There were 220 trials after 20 null trials on day 1 and 40 non-visual trials on day 2. The one-minute break was given every 10 trials to remove short-term memory and encourage long-term memory (Hadjiosif et al., 2023).

### Gas exchange

We recorded the O2-CO2 gas exchange using a portable metabolic measurement system in the breath-by-breath mode (COSMED K5, COSMED Srl, Italy). In Experiment 1, we started recording the RER from the familiarization phase to the end of the experiment to examine how the baseline RER in the pre-experiment, as well as training the RER in the experiment, can predict the fast and slow components of sensorimotor learning and how the RER alters during sensorimotor learning. In Experiment 2, to investigate how glucose intake modulates the RER, we recorded before and after the intervention phase (pre- and post-intervention phases) on day 1. Both phases took 20 min, as determined by the data analysis in Experiment 1, and the data reliability, calculated by the mean squared error against the mean of the whole data, was saturated at 20 min. The gas exchange was recorded continuously during the training phase. On day 2, we recorded the pre-experiment baseline gas exchange for 20 min and then continuously recorded it during the retention phase. Before the experiment, the K5 system was calibrated according to the manufacturer’s instructions.

### Electromyography

In Experiment 1, muscle activity was recorded using a wireless surface electromyography system (WavePlus EMG system, Cometa, Italy). The wireless system had a very low and consistent delay - 14 ms reported by the product specifications - which had little to no impact on the results in reaching movements that took 1980 ± 87 ms. Data were recorded from four muscles: the pectoralis major, posterior deltoid, biceps brachii, and triceps brachii, which govern large portions of shoulder extension/flexion and elbow extension/flexion during horizontal reaching movements. An electrode size was 41 × 15.5 × 11.3 mm and the inter-electrode distance was 20 mm (Pico EMG, Cometa, Italy). The electrode sites were carefully determined by visually checking the data quality during the ballistic reaching movements before the experiment. The EMG and kinematic data were recorded using an A/D signal converter (PCI-6229, National Instruments, USA) at a sampling rate of 1000 Hz. The EMG signals were amplified (×1000) and band-pass filtered from 10 to 500 Hz via the hardware.

### Glucose administration

In Experiment 2, we examined the effects of glucose intake on RER and 24-h retention. We asked the participants in both the WG and Wa groups not to consume any food or drink with sweeteners for at least 1 h before the experiment, to maximize the effects of the intervention. The WG group consumed water and gel-type drinks (Glucorescue, Arkray, Japan), while the Wa group consumed only water during the intervention phase; therefore, they had to take off their masks to record gas exchange. The main ingredient of the gel-type drinks was 50 g of glucose, corresponding to 200 kcal. Because the WG group took 387 ± 40 seconds, the Wa group also waited for approximately 360 seconds, to match the periods of the intervention phase.

## Acknowledgements

We thank Ryota Furuno, On Kobayashi, Yuta Nozawa, Kotaro Beppu, and Asako Munakata for their help in organizing and conducting the experiments. We thank Daichi Nozaki and Maurice Smith for helpful discussions. This work was supported by KAKENHI grants from JSPS (JP23H03296) and JST/PRESTO (JPMJPR23S8) to TH. This work was also partly supported by JST Moonshot R&D Grant Number JPMJMS2239.

## Data analysis

We smoothed the handle position using a fourth-order Butterworth filter with a cutoff frequency of 10 Hz. Subsequently, we found the midpoints of the forward movement, that is, 10 cm away from the start position. This is almost identical to the position at peak velocity, which represents the feedforward predictive component of movement. We computed the movement directions at the midpoints relative to the target direction, because VMR is an angular perturbation. We performed outlier removal in each trial of each experiment if the data were outside the median ± 3 IQRs across participants, which was nearly 0.05% under the hypothetical standard distribution. We removed one participant from Experiment 2 because the data were very distant from the others, especially in the retention phase (38 out of 40 retention trials were outliers).

We used a two-state model to represent short- and long-term memory in sensorimotor learning(Smith et al., 2006). The model considered that the total motor outputs (*x*) were determined by the sum of two distinct memories (*x* = *x*_*f*_+*x*_*s*_) and that the memories are parallelly updated with the error (*e*= *x*+*p, p* is VMR) as *x*_*f*_ (*n*+1)= *α*_*f*_ *x*_*f*_ (*n*)+*β*_*f*_ *e*(*n*) and *x*_*s*_(*n*+1)= *α*_*s*_*x*_*s*_(*n*)+*β*_*s*_*e*(*n*), where *f* and *s* indicate labels of fast and slow components of sensorimotor learning, *n* indicates the trial number, and *α* and *β* are retention and learning factors. The retention and learning factors are thought to be different between the components: 0 < *α*_*f*_ < *α*_*s*_ < 1 and 0 < *β*_*s*_ < *β*_*f*_. We fitted the model to individual data with expectation maximization methods(Albert and Shadmehr, 2018), which allowed us to robustly estimate the parameters of the model from the noisy data; thus, we did not have to obtain a number of samples. A previous report showed that almost 20-95% fewer participants were needed from the power analysis compared to the least-squares algorithm (Albert and Shadmehr, 2018). In Experiment 1, we recruited a number of participants comparable to those in previous studies (Albert and Shadmehr, 2018). Basically, we utilized the original code that is available at http://shadmehrlab.org/tools, although we fixed a few initial parameters for the parameter searching: narrower ranges of initial fast and slow motor memories from ±30° to ±5°. Narrower ranges that emphasized zero learning amounts at baseline were more reflective of the current situation.

The O2-CO2 gas exchange ratio was used to compute the RER offline for each breath. The RER was recorded breath-by-breath and it was obtained at discrete times over a few seconds. For the RER during tasks, we performed linear interpolation between the RER to match the kinematic data with a sampling rate of 1000 Hz and used the RER at the midpoint of movement as a representative index in the trial. The range of RER was theoretically defined as 0.7-1.0, the outsides of which were regarded as outliers. In particular, we removed two participants in Experiment 2 because about 35% of trials were determined as outliers, which is a high ratio compared with the others (7.5 ± 1.0%). For the steady-state RER, such as the pre-experiment phase in Experiment 1 and the pre- and post-intervention phases in Experiment 2, we computed the mean of the RER from the individual breathing data after outlier removal in each breathing data.

EMG data were subtracted from the mean and rectified for each trial. Here, we used muscle activity to estimate the intensity of movement execution, as it affects the RER(Goedecke et al., 2000). Therefore, instead of EMG activity during the middle movement, we computed a trial-wide summation of the EMG of each muscle. We then performed self-normalization with the mean across trials for each muscle. This allowed us to examine how muscle activity changes with sensorimotor learning, independent of the different EMG responses of the muscles.

To examine whether the intensity of the reaching movements influenced the subsequent RER in Experiment 1, we computed a linear regression from muscle activity to RER with a lag within blocks. There were four muscle activities [900 × 4] and RER [900 × 1] over 900 trials. Seventy-five trials were performed in each of 12 blocks; therefore, we cut the blocks for both activities [75 × 12 × 4] and RER [75 × 12 × 1], and slid them every single trial within blocks, as in the preceding trials [1, …, 75-i] and the RER with the i-th lag [i-75]. From the regression results, we computed the coefficient of determination (R^2^), which indicates the percentage of variance explained. For comparison, we randomly permuted the blocks 5000 times, performed the same procedures as described above, and obtained the mean R^2^ across the iterations. We performed this for each participant and then tested the differences in R^2^ between the experimental and permuted data with each of the 75 lags. To compensate for the 75 repetitive tests, we corrected the p-values using the FDR.

For the estimates of intra-individual enhancements in the 24-h retention in Experiment 2, we computed the changes from baseline RER to intervention RER over the regression line (*y*= *ax*+*b*), indicating how much glucose intake increased next-day retention along with the RER changes. Using the slopes (*a*) and offset (*b*) of the regression line, we first predicted the retention levels from the baseline RER (*y*_*pre*_= *ax*_*pre*_+*b*), which indicates the retention of the original RER. We then projected the RER during the task on day 1 onto the regression line (*y*_*task*_= *ax*_*task*_+ *b*) and calculated the extent to which the RER increased within individuals as follows: (*y*_*task*_− *y*_*pre*_)/*y*_*pre*_ × 100 (×). Thus, this value estimates the intra-individual changes in retention due to glucose intake. For comparison, we performed the same procedure in the Wa group.

We performed one-/paired-sample or two-sample two-tailed t-tests to compare the mean variables and performed Pearson’s correlation when testing the covering variables. The significant p-value was set at 0.05.

## Data and code availability

We prepare an online repository on Zenodo (https://zenodo.org/) that contains the data and codes in the current study. The uploads will be completed when the paper is published.

